# Memory B cells and memory T cells induced by SARS-CoV-2 booster vaccination or infection show different dynamics and efficacy to the Omicron variant

**DOI:** 10.1101/2022.07.31.500554

**Authors:** Setsuko Mise-Omata, Mari Ikeda, Masaru Takeshita, Yoshifumi Uwamino, Masatoshi Wakui, Tomoko Arai, Ayumi Yoshifuji, Kensaku Murano, Haruhiko Siomi, Kensuke Nakagawara, Masaki Ohyagi, Makoto Ando, Naoki Hasegawa, Hideyuki Saya, Mitsuru Murata, Koichi Fukunaga, Ho Namkoong, Xiuyuan Lu, Sho Yamasaki, Akihiko Yoshimura

## Abstract

Although BNT162b2 vaccination was shown to prevent infection and reduce COVID-19 severity, and the persistence of immunological memory generated by the vaccination has not been well elucidated. We evaluated memory B and T cell responses to the SARS-CoV-2 spike protein before and after the third BNT162b2 booster. Although the antibody titer against the spike receptor-binding domain (RBD) decreased significantly 8 months after the second vaccination, the number of memory B cells continued to increase, while the number of memory T cells decreased slowly. Memory B and T cells from unvaccinated infected patients showed similar kinetics. After the third vaccination, the antibody titer increased to the level of the second vaccination, and memory B cells increased at significantly higher levels before the booster, while memory T cells recovered close to the second vaccination levels. In memory T cells, the frequency of CXCR5^+^CXCR3^+^CCR6^-^ cTfh1 was positively correlated with RBD-specific antibody-secreting B cells. Furthermore, T cell-dependent antibody production from reactivated memory B cells *in vitro* was correlated to the Tfh-like cytokine levels. For the response to variant RBDs, although 60%-80% of memory B cells could bind to the Omicron RBD, their binding affinity was low, while memory T cells show an equal response to the Omicron spike. Thus, the persistent presence of memory B and T cells will quickly upregulate antibody production and T cell responses after Omicron strain infection, which prevents severe illness and death due to COVID-19.

## Introduction

In December 2019, an outbreak of unknown etiology with apparently viral pneumonia emerged in Wuhan, China. On January 9, 2020, the World Health Organization (WHO) announced the discovery of a novel coronavirus, severe acute respiratory syndrome coronavirus 2 (SARS-CoV-2), which is the pathogen responsible for coronavirus disease 2019 (COVID-19). The COVID-19 pandemic has spread globally, infecting more than 500 million people and causing more than 6.5 million deaths. However, since the development and widespread administration of mRNA vaccines BNT162b2 (Pfizer) and mRNA-1273 (Moderna) encoding the SARS-CoV-2 spike protein as well as the emergence of the Omicron strain, the fatality rate, which was estimated to be about 3% in early 2020, has decreased to less than 0.3% currently, and severe cases have been greatly reduced (1, 2).

Vaccination against SARS-CoV-2 has shown a high preventive effect worldwide, especially for reducing severe illness and death. This vaccination induces a robust antibody response, particularly the emergence of neutralizing antibodies (3–5). However, serum antibody levels drop markedly several months after SARS-CoV-2 infection and vaccination (6). Conversely, persistence and the significance of immunological memory have not been fully investigated.

Mutations frequently occur in the spike glycoprotein in SARS-CoV-2 variants, which can alter viral transmission and immune recognition (7, 8). Omicron variants have amino acid mutations that are concentrated in the receptor binding domain (RBD) of the spike protein, and there is a significant concern that the initial vaccines based on the Wuhan strain may be less effective in preventing infection. However, vaccination could establish immunological memory in B and T cells (5, 9, 10), and it is expected that the third or fourth booster vaccination will strongly reduce COVID-19 severity. However, immune responses after SARS-CoV-2 vaccination vary widely among individuals, and the interaction between humoral and T-cell immunity in defense against COVID-19 have not been fully elucidated.

The humoral response by B cells, specifically antibody production and class switching, is closely regulated by follicular helper T (Tfh) cells. Tfh cells play a central role in facilitating germinal center reactions and promoting differentiation of cognate B cells for antibody secretion (9). CXCR5^+^ circulating Tfh (cTfh) cell are present in human blood, and subdivided into cTfh1, cTfh2, cTfh17, and cTfh1/17 subsets (11, 12). Although cTfh subsets have been reported to be associated with neutralizing antibody titers in COVID-19 patients (13, 14), the relationship between cTfh and humoral responses in vaccinated subjects has not been reported.

In the present study, we comprehensively examined T and B cell immunological memory before and after the third BNT162b2 vaccine booster injection in healthy Japanese volunteers. B cell memory was assessed using FACS for surface expression levels of anti-RBD IgG antibodies, and the ELISpot assay was used to measure the number of B cell that produce anti-RBD antibodies. T cell memory was also examined using FACS analysis of surface activation markers and ELISpot assay for IFNγ after culturing PBMCs with spike protein peptide pools. We investigated memory B and T cell interactions using several approaches. We also evaluated the effects of vaccination from various angles, including cross-reactivity of memory cells to Omicron variants.

## Materials and methods

### Study approval

This study was approved by the Ethics Committee at our University School of Medicine (20200063) and conducted in compliance with the tenets of the Declaration of Helsinki. Informed consent was obtained from all participating individuals.

### Healthy volunteers and COVID-19 convalescent patients

Forty-three healthy individuals from Keio University and Hospital, Tokyo, Japan were enrolled, and their demographic information is shown in Table 1. All healthy individuals had no known history of any significant systemic diseases, including autoimmune disease, diabetes, kidney or liver disease, or malignancy. They were vaccinated three times with a 3-week interval between the first and second vaccinations and an 8-month interval between the second and third vaccinations. Eighty-eight patients who had COVID-19, which was diagnosed using approved RT-PCR tests for SARS-CoV-2 with nose swabs or saliva, and who were hospitalized at Keio University Hospital between April and December 2020 were enrolled (15). Patient information is listed in Table 2 and Supplemental Table 1. Blood samples from patients were collected at outpatient visits over 6 months to 1 year.

**Table 1.** Demographic characteristics of vaccinated healthy volunteers (n= 43)

| Age | Sex |  | Sample collection Date |
| --- | --- | --- | --- |
|  | Female | Male |  |
| 28 ~ 62 | 55.8%<br>(24/43) | 44.2%<br>(19/43) | March 2021 ~ January 2022 |

**Table 2.** Demographic and clinical characteristics of SARS-Cov-2 recovered patients (n = 88)

| COVID-19 severity |  | Female | Male | Number<br>of<br>sampling | Sample collection<br>days<br>after PCR positive | Age | Pre-existing disease |
| --- | --- | --- | --- | --- | --- | --- | --- |
| asymptomatic | 16<sup>a</sup><br>(18.2 %)<sup>b</sup> | 6<br>(37.5 %) | 10<br>(62.5 %) | 1 | 25 ~ 84 | 23 ~ 62<br>(34.9)<sup>c</sup> | 9<br>(56.3 %) |
| mild | 22<br>(25.0 %) | 11<br>(50.0 %) | 11<br>(50.0 %) | 1 ~ 3 | 12 ~ 202 | 22 ~ 58<br>(32.5) | 15<br>(68.2 %) |
| moderate | 39<br>(44.3 %) | 13<br>(33.3 %) | 26<br>(67.7 %) | 1 ~ 3 | 10 ~ 221 | 20 ~ 78<br>(50.1) | 31<br>(79.5 %) |
| severe | 5<br>(5.7 %) | 1<br>(20.0 %) | 4<br>(80.0 %) | 1 ~ 3 | 33 ~ 208 | 47 ~ 62<br>(58.8) | 5<br>(100 %) |
| critical | 6<br>(6.8 %) | 0<br>(0 %) | 6<br>(100 %) | 1 ~ 3 | 39 ~ 242 | 52 ~ 74<br>(62.0) | 5<br>(83.3 %) |

<sup>a</sup> The number of patients is indicated. <sup>b</sup> The percentage of patients is indicated. <sup>c</sup> The average of age is indicated.

### Isolation of human PBMCs

Blood was diluted with PBS and then gently loaded onto the Lymphoprep^TM^ (Serumwerk Bernburg AG, Bernburg, German) layer with a density of 1.007 ± 0.001 g/mL (20°C) followed by density gradient centrifugation (820 ×*g*, 25°C, 30 min). Plasma samples were aliquoted and stored at −20°C after density gradient centrifugation. Cells were washed with PBS containing 0.5% BSA and 2 mM EDTA, and then cryopreserved in CellBanker 1plus (TAKARA Bio, Kusatsu, Japan) at –80°C until use.

### SARS-CoV-2 spike protein RBD production

Whole spike protein and the spike protein RBD were prepared as described previously (16, 17). Briefly, the RBD was inserted into pcDNA3.4 with a streptavidin binding peptide (SBP) tag at the C-terminus and were produced using the Expi293 Expression System (Thermo Fisher Scientific, Waltham, MA) according to manufacturer’s instruction. The supernatant containing the RBD was collected and purified using Streptavidin Sepharose High Performance beads (Cytiva, Tokyo, Japan). The protein purity was determined by SDS-PAGE, and the concentration was determined using a bicinchoninic acid (BCA) Protein Assay Kit (ThermoFisher Scientific). Biotinylated SARS-CoV-2 spike RBD, His, Avitag^TM^ of both Wuhan and Omicron types were purchased from ACROBiosystems (Newark, DE).

### ELISA

The RBD that we made was diluted to 5 μg/mL in PBS and coated onto flat 96-well plates (442404, ThermoFisher Scientific) overnight. When using commercially sourced RBD, 10 μg/ml of streptavidin (S4762, Sigma-Aldrich, St Louis, MO) was coated overnight and RBD at the concentration of 0.5 μg/ml was applied on streptavidin-coated plates after washing with PBS. The plates were blocked with blocking buffer (Bethyl Laboratories, Montgomery, TX) for 30 min at room temperature (RT). After washing with PBS containing 0.05% Tween-20, plates were incubated with plasma diluted 1:1000-5000 using diluent buffer (blocking buffer with 0.05% Tween-20) for 2 h at RT. After washing, plates were incubated with Peroxidase AffiniPure F(ab[)[Fragment Goat Anti-Human IgG (Jackson Immunol Research, West Grove, PA) at a 1:5000 dilution for 1 h. After the final washing, plates were incubated with TMB Substrate Set (BioLegend, San Diego, CA) for 10 min. Reactions were stopped with H_2_SO_4_, and the optical density at 450 nm (OD450) was measured. Recombinant human SARS-CoV-2 spike S1 IgG lyophilized antibody (Clone AM009105, BioLegend) was used as a standard.

### ELISpot assay of IFN-γ producing cells

Fifteen mer peptide pools for SARS-CoV-2, PepTivator SARS-CoV-2 Prot_S (130-126-700), S1 (130-127-041), and S+ (130-127-311) were obtained from Miltenyi Biotec (Bergisch Gladbach, Germany) and mixed equally to use for stimulation.

As peptide pools derived from the Omicron variant, the PepTivator® SARS-CoV-2 Prot_S B.1.1.529 Mutation Pool (130-129-928, Miltenyi Biotec) and SARS-CoV-2 Spike Glycoprotein B.1.1.529-Omicron (RP30121, GenScript Biotecn Corp, Piscataway, NJ) were used with the simultaneous control of the wild-type, PepTivator® SARS-CoV-2 Prot_S B.1.1.529 WT Reference Pool (130-129-927, Miltenyi Biotec) and SARS-CoV-2 Spike Glycoprotein-crud (RP30020, GenScript Biotecn Corp). A ten amino acid overlapping peptide mixture pool was prepared, the details of which will be published elsewhere.

A human IFNγ ELISpot^plus^ kit (MABTECH AB, Nacka Strand, Sweden) was used in accordance with the manufacturer’s instructions. PBMCs were thawed and washed with culture medium (RPMI-1640 containing 10% FBS, 2 mM L-glutamine, 1 mM sodium pyruvate, non-essential amino acids, 10 mM Hepes, penicillin-streptomycin, and 2-mercaptoethanol). After incubation with culture medium for 30 min, 3×10^5^ cells were added to each well in the ELISpot plates, which were blocked with culture medium, with or without 60 nM of mixtures of peptide pools. In some experiments, PBMCs were cultured with oligopeptide mixtures for 1–2 days, and then cells were applied into ELISpot plate wells. The spots were tested using detection antibody and Streptavidin-ALP. Spots were read using a CTL-ImmunoSpot S5 Series Analyzer (Cellular Technology Limited, Shaker Heights, OH).

### ELISpot assay of anti-RBD antibody-producing cells

For polyclonal activation, 2 × 10^5^ PBMCs were cultured for 4 days with 1 μg/mL R-848 (Resiquimod, MedChem Express, Monmouth Junction, NJ), an agonist of TLR7/8, in the presence of 5 ng/mL rhIL-2 (Peprotech, Cranbury, NJ) and applied onto the ELISpot plate using the Human IgG Single-Color ELISPOT Assay (Cellular Technology Limited), which was coated with anti-human Igκ and anti-human Igλ capture antibodies and then blocked as described in the manufacturer’s instructions. After overnight culture, cells were washed out and 0.5 μg/mL SBP-tagged RBD or biotinylated RBD was added. To detect total IgG-secreting cells, anti-human IgG detection antibody was used. Anti-RBD antibody among the total IgG was detected using Streptavidin-ALP and substrates in the kit.

### FACS analysis of AIM^+^ cTFh subsets and RBD-MBCs

The activation-induced markers (AIM) assay was performed as previously described (18). PBMCs were cultured in the presence of 60 nM peptides or 1 μg/mL recombinant spike protein for 1–2 days, and cells were stained with CD4, CD8, CD69, CD137, and OX40 antibodies. For memory and cTfh analysis, CD45RA, CCR7, CXCR5, CXCR3, and CCR6 were stained as previously described (19). cTfh cells were divided into cTfh1 (CXCR5^+^CXCR3^+^CCR6^-^), cTfh2 (CXCR5^+^CXCR3^-^CCR6^-^), and cTfh17 (CXCR5^+^CXCR3^-^CCR6^+^) subsets.

To detect anti-RBD IgG antibody-expressing B cells, the thawed PBMCs were incubated with 10 μg/mL SBP-tagged RBD or 1 μg/mL biotinylated RBD plus FcR Blocking Reagent human (Militenyi Biotech) in PBS containing 1% FBS, 2 mM EDTA, and 0.04% NaN_3_ for 30 min at 37°C. After washing, the cells were stained with mixture of streptavidin-Brilliant Violet^TM^ (BV)-410 (Biolegend) and strepatavidin-PE/Cy7 (Invitrogen), and monoclonal antibodies against CD20, CD3, CD11b, IgG, CD19, and CD27, and fixable viability dye efluor™ 780 (ThermoFisher Scientific). The clones and antibody sources are listed in Supplemental Table 2. Stained samples were applied on FACS Canto II (BD, Franklin Lakes, NJ) or Beckman Coulter CytoFlex S (Beckman Coulter, Brea CA), and analyzed using FlowJo10.5.3 (Sheffield, UK).

### In vitro antibody production assay

PBMCs (3 × 10^5^ cells) were cultured with 60 nM of a 15 amino acid oligopeptide mixture or 1 μg/mL recombinant whole spike protein for 4 days in the presence of 5 ng/mL rhIL-2 in 96-well round bottom plates and then washed with culture medium twice. Culture was continued for another 3 days without oligopeptide or spike protein. Culture supernatant was harvest and the anti-RBD antibody concentration was measured using ELISA.

### Quantitative PCR

PBMCs were cultured for 24 hours as described above, and total RNA was purified using the ReliaPrep™ RNA Miniprep System (Promega, Madison, WI). cDNA was obtained using a High-Capacity cDNA Reverse Transcription Kit (Applied Biosystem Waltham, MA). Real-time PCR was performed by CFX Connect Real-Time PCR Detection System (Bio-Rad, Hercules, CA) with SsoAdvanced Universal SYBR Green SuperMix (Bio-Rad). The following primers were used: hHPRT-f, TGAGGATTTGGAAAGGGTGT; hHPRT-r, CCTCCCATCTCCTTCATCAC; hIL21-f, GTGAATGACTTGGTCCCTGAA; hIL21-r, AAGCAGGAAAAAGCTGACCA; hIL4-f, TCAAAACTTTGAACAGCCTCA; hIL4-r, CTTGGAGGCAGCAAAGATGT; hIFNG-f, TGCCAGGACCCATATGTAAA; hIFNG-r, TCCATTATCCGCTACATCTGAA; hIL10-f, CTGGGGGAGAACCTGAAGA; and hIL10-r, GGCCTTGCTCTTGTTTTCAC.

### Statistical analysis

The numerical data were statistically analyzed and visualized using GraphPad Prism 9 software (GraphPad Software, San Diego, CA). Multiple data were analyzed with Kruskal–Wallis test and followed by Dunn’s multiple comparisons test. Unpaired data were analyzed using the Mann–Whitney test. Differences with P values less than 0.05 were considered to be significant: \**p* < 0.05, \*\**p* < 0.01, \*\*\**p* < 0.001, \*\*\*\**p* < 0.0001, and ns (not significant for *p* ≥ 0.05). The relationship between variables was studied using a simple linear regression model in the software R (https://www.R-project.org/).

## Results

### Antibodies in plasma rapidly decreases, while memory B cells progressively increase after vaccination

Serum antibody levels and B cell memory were evaluated from 43 volunteers listed in Table 1 at the following three time points: 3 weeks after the second vaccination (V2_3w); 8 months after the second vaccination (V2_8m); and 3 weeks after the third vaccination (V3_3w). As reported (6, 20), plasma anti-RBD antibody levels drastically increased at V2_3w compared with those of pre-vaccination and then decreased to almost undetectable levels at V2_8m. Approximately 3 weeks after the third vaccination (V3_3w), antibody levels recovered to similar or even higher levels compared with those at V2_3w (Fig. 1A and Supplemental Fig. 1A). To measure RBD-specific memory B cells, we established the following two methods: FACS analysis and ELISpot assay using purified RBD containing an SBP tag (17). As shown in Fig. 1B and Supplemental Fig. 1B, RBD-specific IgG-expressing B cells were detected with two different RBD-fluorochromes after CD19^+^ and IgG^+^ gating. RBD-specific IgG-expressing B cells could be detected at V2_3w (0.263 ± 0.197% in CD19^+^IgG^+^ cells), and increased up to 8 months after the second vaccination (0.535 ± 0.281%, Fig. 1B–C, Supplemental Fig. 1C). Because most of the RBD-specific IgG-expressing B cells exhibited CD20^high^ (Fig. 1D) memory phenotypes, we confirmed them as RBD-specific memory B cells (RBD-MBCs). RBD-MBCs further increased after the third vaccination (V3_3w) (1.759 ± 1.055%).

**FIGURE 1.**
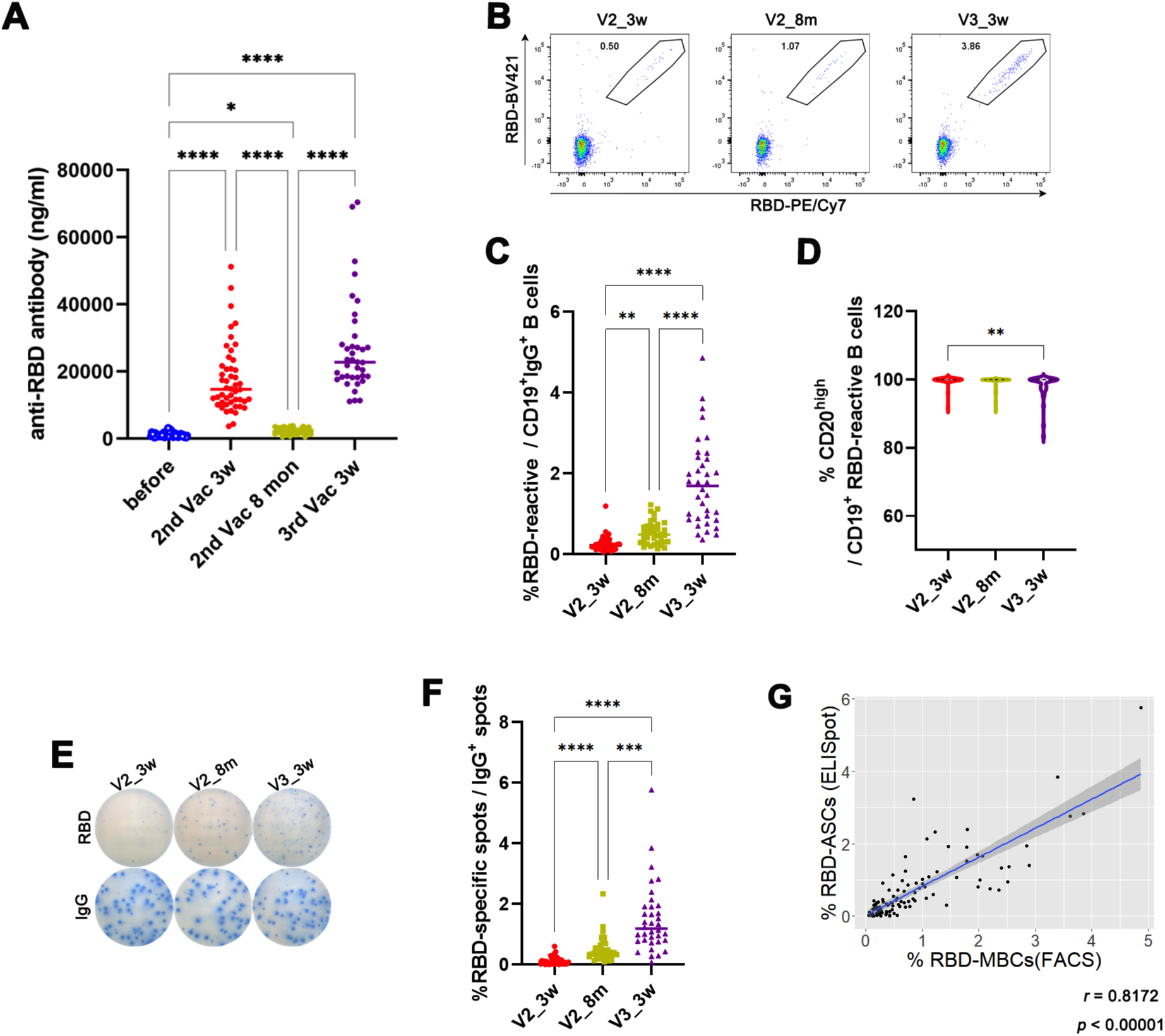
Anti-RBD antibody concentration in plasma decreased, but the MBC was continuously increased until 8 months after vaccination. **(A)** The anti-RBD-antibody concentration in plasma of healthy volunteers after 3 weeks (V2_3w) and 8 months (V2_8m) after the second vaccination and at 3 weeks after the third vaccination (V3_3w). **(B)** Representative FACS profiles of RBD-MBCs at the indicated time points. Gating strategy is indicated in Supplemental Fig. 1B. **(C)** RBD-MBCs frequencies. **(D)** The percentages of CD20^high^ population in RBD-MBCs. **(E)** Spots indicate RBD-ASCs in 2 x 10^5^ PBMCs and IgG secreting B cells in 2 x10^3^ cells. **(F)** The percentages of RBD-ASCs at each time point are indicated. **(G)** A positive correlation exists between RBD-MBCs and RBD-ASCs. The Kruskal–Wallis test was performed, and *p* values were determined using Dunn’s multiple comparisons test. *, *p* < 0.05; **, *p* < 0.01; ***, *p* < 0.001; ****, *p* < 0.0001.

Next, we evaluated anti-RBD antibody-secreting B cells by the ELISpot assay. To facilitate differentiation of memory B cells to antibody-secreting plasma cells, PBMCs were cultured with R848, a potent agonist of TLR7/8 (21, 22). Without R848 stimulation, no or few spots appeared, indicating that most spots were derived from the antibody from plasma cells that were converted from memory B cells in the blood (data not shown). The number of RBD-specific antibody-secreting B cells (RBD-ASCs) detected using a RBD-SBP tag was counted and normalized by dividing by the total number of IgG-secreting B cells detected using anti-IgG antibody (Fig. 1E–F). Consistent with RBD-MBCs detected by FACS, RBD-ASCs were detected at V2_3w (0.137 ± 1.139% per IgG secreting cells), progressively increased at V2_8m (0.510 ± 0.408%), and profoundly enhanced by the third vaccination (1.489 ± 1.114%, Fig. 1F and Supplemental Fig. 1D). The frequencies of RBD-MBCs and RBD-ASCs were well correlated (Pearson’s correlation *r=* 0.817; Fig. 1G). Consistent with other reports (9, 10), these data indicate that although the antibody levels against SARS-CoV-2 had severely decreased 8 months after the second vaccination, memory B cells were circulating in the blood, and they were progressively increased by the booster vaccination.

### T cell memory maintained after vaccination and cTfh subset was correlated with B cell response

Next, to evaluate T cell response, IFNγ-ELISpot assay and AIM^+^ FACS assay were performed after 1–2 days of PBMC stimulation with 15 mer or 10 mer peptide pools of the SARS-CoV-2 spike protein (23). Fig. 2A shows representative IFNγ spots that were detected after 1 day or 2 days of culture that appeared at V2_3w, decreased at V2_8m to about half the level, and then were restored after the third vaccination (Fig. 2B, Supplemental Fig. 2A, for 1 day culture and Supplemental Fig. 2B for 2 days culture). The activation markers OX40 and CD137 were also examined using the AIM assay after culture with the 15 mer peptide pools. Activation markers were upregulated in CD4^+^ T cells but there was minimal upregulation in CD8^+^ T cells (Fig. 2C-D), suggesting that 15 mer oligopeptides stimulated mostly CD4^+^ T cells. On the other hand, 10 mer peptide pools induced IFNγ expression from CD8^+^ T cells but not from CD4^+^ T cells after 2 days of culture (Supplemental Fig. 2C), indicating that 10 mer peptides are more suitable than 15 mer peptides for detection of antigen-specific CD8^+^ T cells. After stimulation with the 10 mer peptide pool, IFNγ spots were also highly maintained regardless of the third booster vaccination (Fig. 2E-F). Thus, we confirmed that both CD4^+^ and CD8^+^ T cell memories were maintained after 8 months with a relatively low decline, and they recovered to the initial levels by the booster vaccination.

**FIGURE 2.**
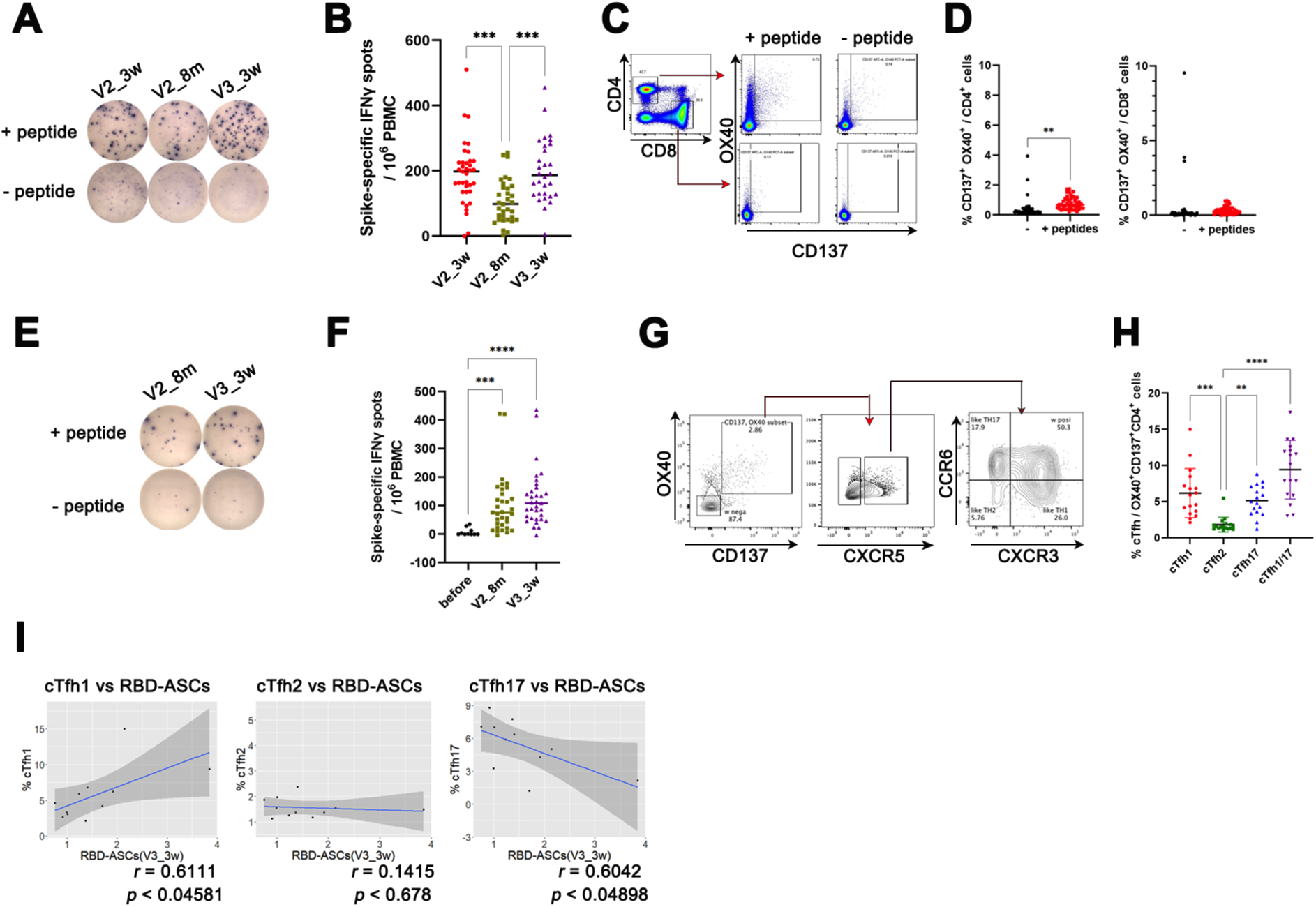
T cell responses against RBD decrease after vaccination, and the early phase of T cell response may affect the subsequent B cell response. **(A)** ELISpot assays of IFNγ after PBMCs stimulation with 15 mer oligopeptide pools of spike protein are shown at the indicated time point. **(B)** The numbers of spots at each time point. **(C)** FACS profiles of OX40^+^CD137^+^ in CD4^+^ or CD8^+^ T cells after 15 mer peptide stimulation. Singlets were identified using FSC and SSC, and then gated on CD3e^+^ and FVD^-^ cells. These cells were separated into CD4^+^ and CD8^+^ cells. **(D)** The percentage of OX40^+^CD137^+^ in CD4^+^ or CD8^+^ T cells after 15 mer peptide stimulation. **, *p* < 0.01 calculated using the Mann–Whitney test. **(E)** ELISpot assays of IFNγ after stimulation of PBMCs with 10 mer oligopeptide pools are shown. (**F**) The numbers of spots are indicated at each time point. (**G**) Representative FACS profiles detected cTfh1, cThf2, cTfh17, and cTfh1/17 after 15 mer peptide stimulation. CD4^+^ cells are described in (**C**) and gating strategy is indicated. **(H)** The percentages of cTfh1, cThf2, cTfh17, and cTfh1/17 at 3 weeks after the second vaccination are shown. (**I**) Pearson’s correlation coefficients are calculated between each cTfh subset at V2_3w and RBD-ASCs at V3_3w. The Kruskal–Wallis test was performed, and *p* values were determined using Dunn’s multiple comparisons test. **, *p* < 0.01; ***, *p* < 0.001; ****, *p* < 0.0001.

cTfh cell subsets were reported to be associated with antibody response in COVID-19 convalescent patients (14, 19, 24, 25). We observed CXCR5^+^ cTfh subsets in OX40^+^ CD137^+^ activated CD4^+^ cells after 15 mer peptide stimulation for 2 days at V2_3w, an early time point in the induction of immune response. As shown in Fig. 2G, cTfh1 (CXCR3^+^CCR6^-^), cTfh17 (CXCR3^-^CCR6^+^), cTfh2 (CXCR3^-^CCR6^-^), and cTfh1/17 (CXCR3^+^CCR6^+^) populations were determined and the frequency of each subset is shown in Fig. 2H. Correlation between these subsets and RBD-ASCs were examined at the V2_3w and V3_3w time points. A positive correlation between the cTfh1 subset and RBD-ASCs at V3_3w (Fig. 2I) was observed but not at V2_3w (Supplemental Fig. 2D). Conversely, cTfh17 was negatively correlated with RBD-ASCs at V3_3w but not at V2_3w. No correlation between cTfh2 and RBD-ASCs was found. Thus, a higher cTfh1 response may result in more memory B cells. These data are consistent with those of a previous report showing a correlation between cTfh1 and B cell memory in case of infection (24).

### In vitro anti-RBD antibody production in PBMC from vaccinated volunteers

Next, we reconstituted the interaction between antibody-producing B cells and CD4^+^ helper T cells *in vitro*. We expected memory B cell activation and differentiation into antibody producing plasma cells by interacting with activated memory CD4^+^T cells. PBMCs were cultured with the peptide pool (S-peptide) or recombinant spike (S-protein) for 4 days in the presence of IL-2, and the culture was continued for another 3 days without peptides or the S-protein. The culture supernatant was removed and the anti-RBD antibody concentration was measured using an ELISA. As shown in Fig. 3A, anti-RBD antibody was detected in several, but not all, PBMCs form vaccinated volunteers. Antibody was more frequently detected in samples that were stimulated with S-peptides than by the S-protein, and only a few PBMC samples produced anti-RBD antibody in response to both stimulations. The most frequent antibody production was observed in PBMCs 3 weeks after the third vaccination (V3_3w), and lowest antibody production was observed at 8 months after the second vaccination (V2_8m), which was only weakly correlated with the memory B cell (Fig. 1C) or memory T cell (Fig. 2B) levels.

**FIGURE 3.**
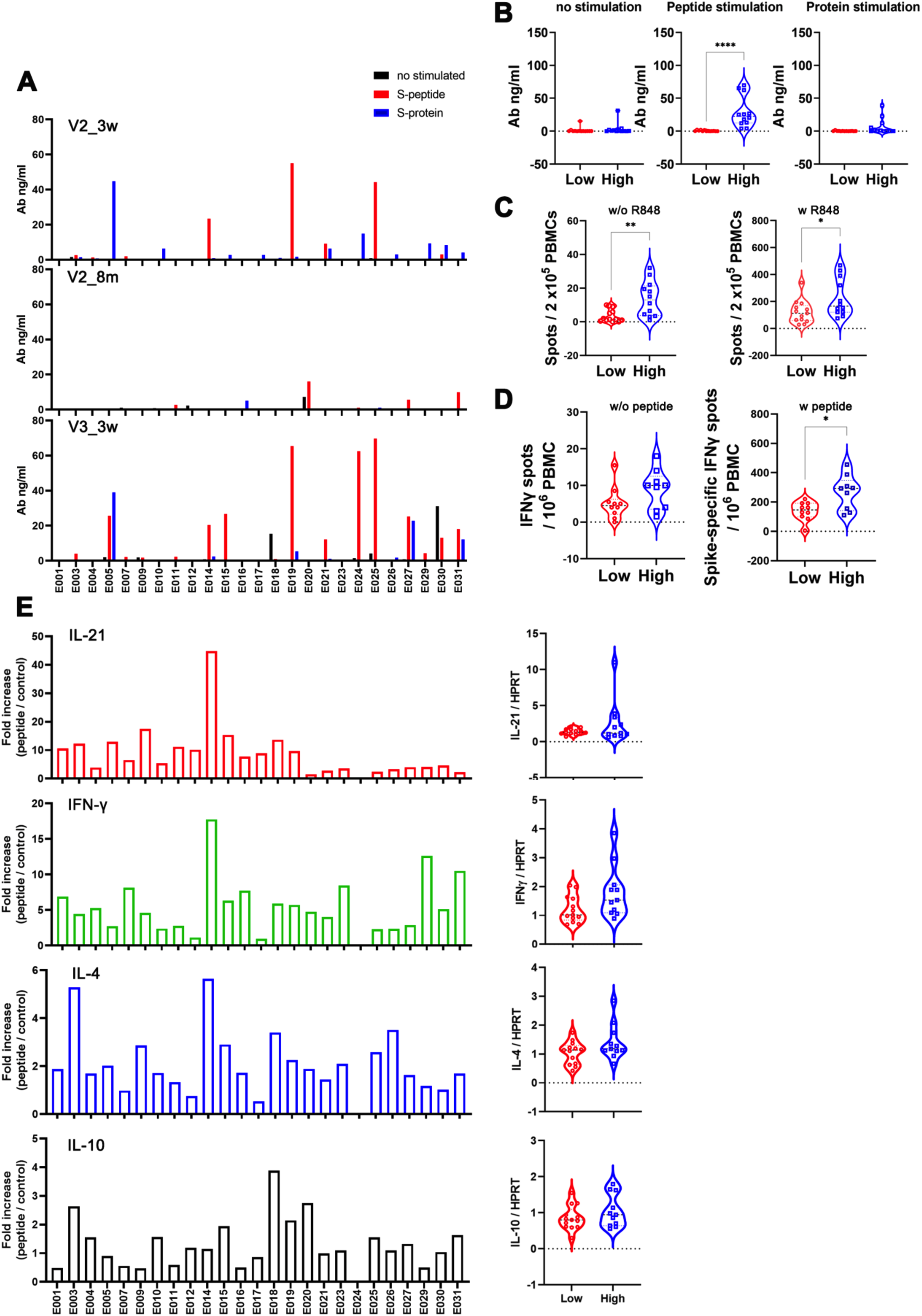
Different anti-RBD antibody production ability after PBMC stimulation *in vitro*. **(A)** *In vitro* antibody production by culturing PBMCs with 15 mer peptides or S protein at the indicated time points. **(B-E)** The V3_3w samples are separated into two groups with low antibody production (E001, E004, E007, E010, E011, E012, E016, E017, E018, E020, E023, and E026) and high antibody production (E003, E005, E014, E015, E019, E021, E024, E025, E027, E029, E030, and E031). (**B**) The antibody concentrations in the culture supernatants in each group are shown after each stimulation. **(C)** The numbers of RBD-ASCs from the ELISpot assay after culture with or without R848 are indicated separately in the low and high antibody production groups. **(D)** The IFNγ-spot numbers from the ELISpot assay after culture with or without the peptide pool are indicated separately in the low and high antibody production groups. (**E**) *IL21*, *IFNG*, *IL4*, and *IL10* mRNA expression in PBMCs cultured with 15 mer peptide pool from each volunteer (left) and summarized in the low and high antibody production groups (right). *, *p* < 0.05; **, *p* < 0.01; ****, *p* < 0.0001 calculated using the Mann–Whitney test.

To clarify the relationship between the *in vitro* antibody-producing ability and memory T and B cells, we separated the samples at V3_3w into two groups, a high antibody-producing group (E003, E005, E014, E015, E019, E021, E024, E025, E027, E029, E030, and E031) and a low antibody-producing group (E001, E004, E007, E010, E011, E012, E016, E017, E018, E020, E023, and E026), and antibody production and T cells and B cell responses were analyzed. Antibodies were highly produced when stimulated with the peptide pool than with the S-protein stimulation in the antibody-producing group (Fig. 3B), suggesting that TCR stimulation of CD4^+^ helper T cells plays a critical role in this culture system. When the numbers of memory B cells (RBD-ASCs) are compared between the two groups, the high antibody group contained slightly more RBD-ASCs than the low antibody group (Fig. 3C). However, the high antibody-producing group exhibited a higher T cell response ability (Fig. 3D). We also examined the expression levels of cytokines related to antibody production in the cultured cells using qPCR (Fig. 3E). Samples in the high antibody group expressed higher *IL21*and *IFNG* mRNA levels. Although induction of *IL4 and IL10* was not as strong as that for *IL21*and *IFNG*, these two cytokine levels were also higher in the high antibody group than in the low antibody group. These data indicate that, in addition to a high B cell memory, T cell responses play important roles in antibody production *in vitro*.

### Memory cells emerged in infected patients with SARS-CoV-2 with similar kinetics as vaccination

We also evaluated memory B and T cells in 88 convalescent COVID-19 patients who had asymptomatic, mild, moderate, severe, or critical symptoms, which are listed in Table 2. Information about each patient is presented in Supplemental Table 1. Although asymptomatic participants provided only one blood sample, most other participants provided blood samples two or three times at various intervals. All severe and critical patients and four of 39 moderate patients received systemic steroid treatment. We measured RBD-specific B cells with FACS (Fig. 4A) and memory T cells with IFNγ-ELISpot assay (Fig. 4B) using samples within 80 days post-infection. Memory B cells were barely detectable in asymptomatic patients. Although memory B cells were detected at the convalescent stage in all mild, moderate, severe, and critical patients, the frequency of RBD-specific B cells was lower than that in vaccinated samples (Fig. 4A and Fig. 1C). RBD-specific T cell memory was detected at the convalescent stage in moderate, severe, and critical patients (Fig. 4B). Although the frequencies of IFNγ-positive memory T cells vary among patients, some of these patients have levels that are comparable to those in vaccinated patients (Fig. 2B). These data suggest that SARS-CoV-2 infection resulted in less efficient memory B cell generation and memory T cell induction at certain levels compared with vaccination.

**FIGURE 4.**
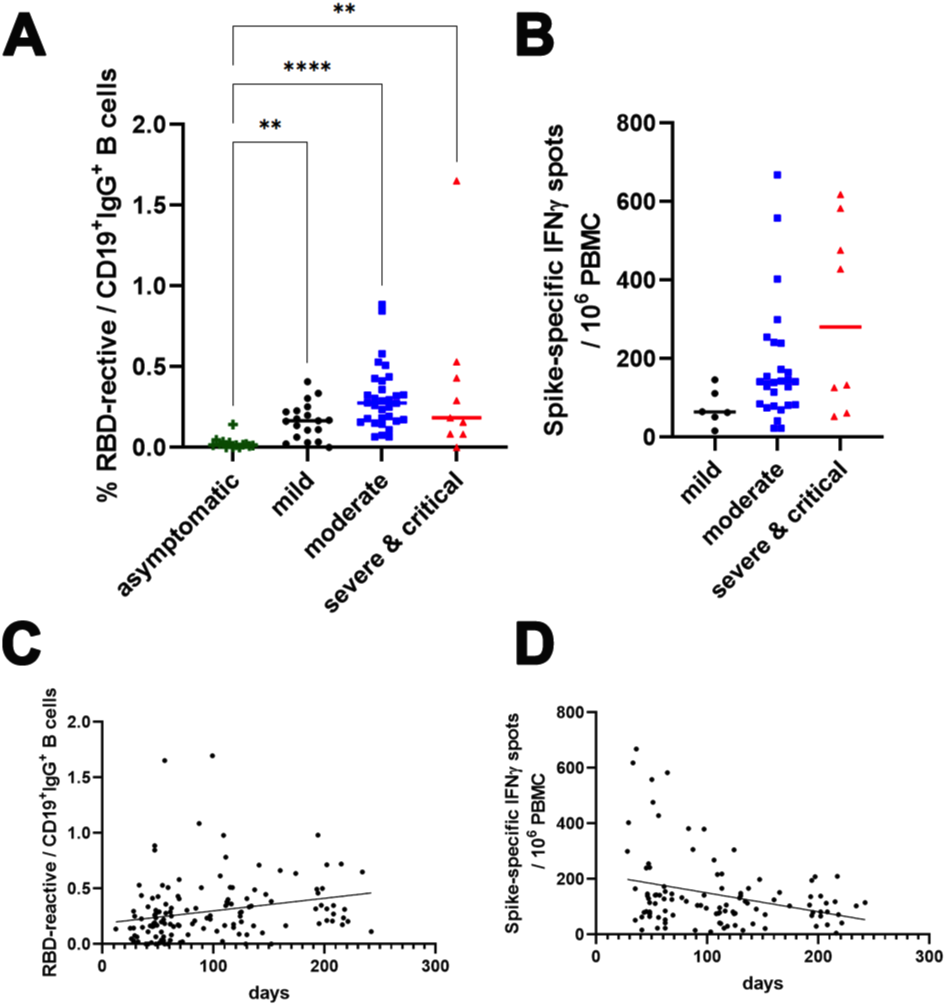
RBD-MBCs and memory T cells in COVID-19 patients are detected. **(A)** The frequencies of RBD-MBCs from the COVID-19 patients within 80 days after a SARS-CoV-2 PCR-positive test result are indicated by COVID-19 severity. **(B)** Spike-reactive memory T cell frequencies from COVID-19 patients within 80 days after a SARS-CoV-2 PCR-positive test result are indicated by COVID-19 severity. **(C)** Cross-sectional analysis of RBD-MBC frequencies detected by FACS as a function of days after a SARS-CoV-2 PCR-positive test result. **(D)** Cross-sectional analysis of memory T cell frequencies as a function of days after a SARS-CoV-2 PCR-positive test result. The Kruskal–Wallis test was performed, and *p* values were determined using Dunn’s multiple comparisons test. **, *p* < 0.01; ****, *p* < 0.0001.

Cross-sectional and longitudinal analysis of RBD-specific memory B cell and T cell frequencies were performed as a function of days after PCR-confirmed infection (Fig. 4C-D for overall patients and Supplementary Fig. 3A-B for patients categorized by symptom severity). Similar to vaccination, RBD-specific memory B cells increased progressively, while memory T cells decreased slowly. Thus, there is less difference in the RBD-specific memory B cell and T cell frequencies among the symptom categories in the samples over 80 days post-infection (Supplemental Fig. 3C-D). These data are consistent with previous reports showing the development and persistence kinetics of memory B and T cells elicited by natural infection with SARS-CoV-2 (23, 24).

### Memory B cells induced by vaccination show lower affinity for Omicron RBD

Vaccination with the parental Wuhan type SARS-CoV-2 has been shown to develop antibodies that can cross-react with other variants, such as Beta and Delta variants (26–29). As expected, the serum antibody titer at V3_3w was significantly lower against the Omicron BA.1 variant RBD, at about 30% of the levels shown in response to the parental Wuhan strain RBD (Fig. 5A). To evaluate whether memory cells are cross-reactive against the Omicron variant, we examined RBD-specific memory T cells using the IFNγ-ELISpot assay (Fig. 5B) and RBD-specific memory B cells using the ELISpot assay (Fig. 5C) and FACS (Fig. 5D). As reported (30, 31), T cell memory responded equally to both the Wuhan and Omicron-type spike peptide pools (Fig. 5B). We used two commercially available peptide pools that contain only mutated peptides or whole spike peptide pools, but both of them showed similar results.

**FIGURE 5.**
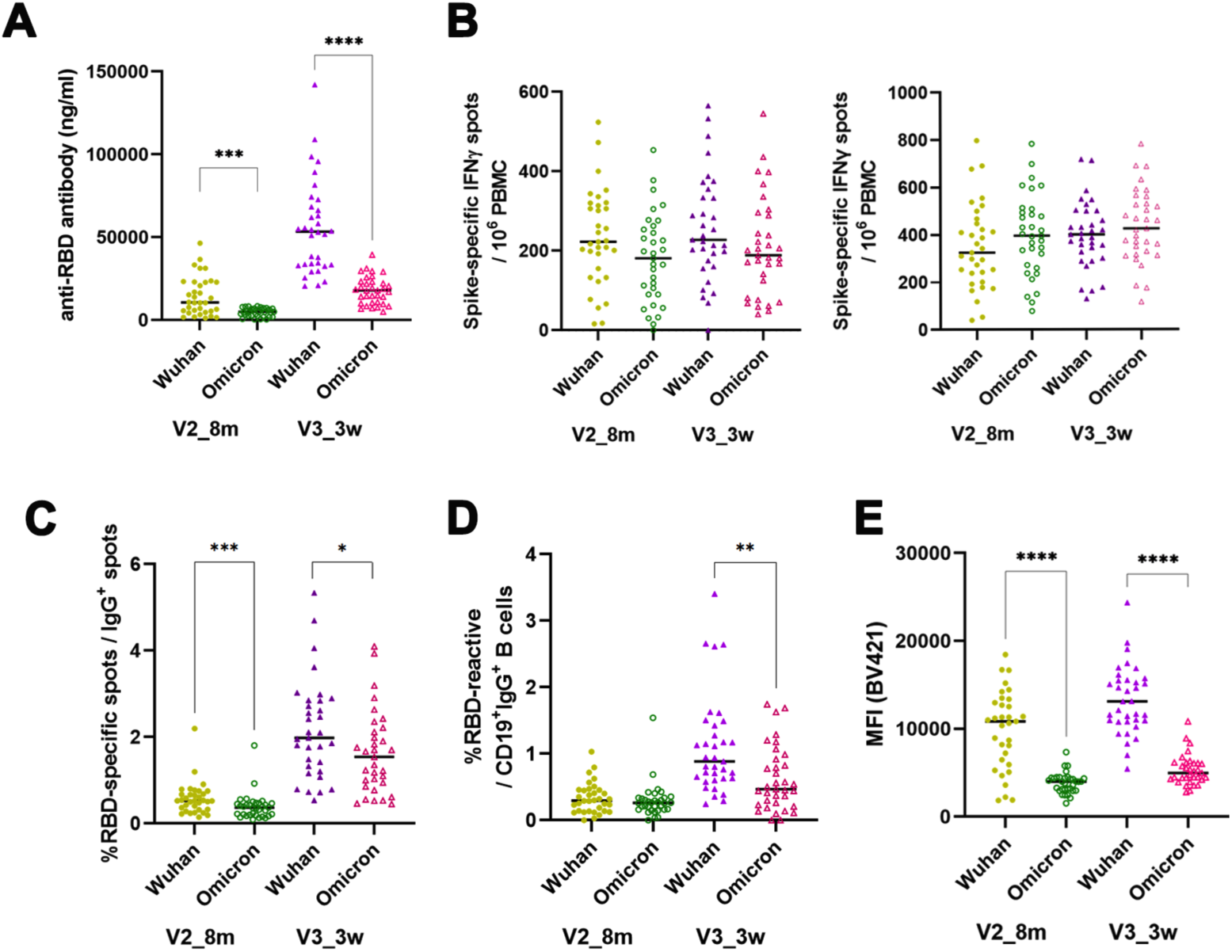
Memory B cells and T cells are both reactive to the SARS-CoV-2 Omicron variant. **(A)** Cross-reactivity of antibody from the plasma of volunteers at V2_8m and V3_3w. **(B)** IFNγ response after 2 days of culture with the Wuhan and Omicron peptide pools of the spike protein derived from two different commercial sources (left, Meliteny Biotec; right, GenScript Biotec Inc.) (**C**) The frequencies of RBD-ASCs reactive to both the parental Wuhan virus and the Omicron variant are shown. **(D)** The frequencies of RBD-MBCs reactive to both the parental Wuhan virus and Omicron variant are shown. **(E)** MFIs of RBD-biotin-SA-BV421 in RBD-MBCs are shown upon reacting to the parental Wuhan virus or the Omicron variant. *, *p* < 0.05; **, *p* < 0.01; ***, *p* < 0.001; ****, *p* < 0.0001 calculated with Mann–Whitney test.

Unexpectedly, the number and frequency of RBD-specific memory B cell memory detected using the ELISpot and FACS assays were not severely reduced against the Omicron RBD (69.4% (V2_8m) and 77.9% (V3_3w), Fig. 5C; and 88.3% (V2_8m) and 56.0% (V3_3w), Fig. 5D). Representative ELISpot data showed that WT and Omicron RBDs did not react with PBMCs collected before vaccination (Supplemental Fig. 4A). As shown by a representative FACS profile (Supplemental Fig. 4B), the mean fluorescence intensity (MFI), which represents the affinity of antibody for RBDs, was lower for Omicron RBD than for the Wuhan type RBD (38.7% (V2_8m) and 40.1% (V3_3w), Fig. 5E). These data indicate that although memory B cells elicited by the BNT162b2 vaccine possess BCRs that are reactive to the Omicron RBD, their affinity was lower than that of the Wuhan type RBD, and therefore, the antibody titer against the Omicron RBD was low (Fig. 5A). Because the T cell response was normal against Omicron, infection with the Omicron variant or vaccination with the Omicron-type spike protein may induce affinity maturation that is high enough to protect against an Omicron infection.

## Discussion

Although it is known that memory B and T cells emerge as a result of BNT162b2 vaccination, the interaction between B and T cells is not well understood. We found that existing antibodies gradually decrease after vaccination and infection, but memory B cells producing anti-RBD antibodies progressively increase in the plasma of vaccinated and infected individuals, which is similar to other reports (9, 28). Memory B cells persisted up to 8 months after the second injection, and they further increased after the third booster. The decrease in serum antibody levels is probably due to a short lifespan of the antibody-producing plasma cells. However, it is not clear why memory B cells continue to increase in the absence of infection after the second vaccination. One possibility is that the amount of spike protein produced by the mRNA vaccine is so high that B cells are still differentiating into plasma cells 3 weeks after the second vaccination, therefore, memory B cells require longer time to accumulate. Alternatively, it is possible that the production of B cell memory is ongoing in the lymph nodes and that it takes time for them to emerge into the blood. As shown recently, memory T and B cells are present in the bone marrow, spleen, lung, and multiple lymph nodes for up to 6 months after infection (32). To verify such possibilities, it would be necessary to measure both the IgM-type memory B cells and the IgG type, and to measure memory B cell levels in the lymph nodes. However, memory B cells persist for a longer time than expected after vaccination; they expand with booster vaccination and they probably also expand with natural Omicron infection. On the other hand, memory T cells that suppress severe disease can respond to both Wuhan and Omicron to the same degree and they live at least more than 8 months. Thus, fourth and fifth booster vaccination is questionable at least for young healthy people who acquired sufficient memory B and T cells by the third vaccination.

Recent studies indicate that Omicron infection to unvaccinated individuals shows the limited neutralization antibody of only Omicron itself, however, infection to vaccinated individuals induces higher neutralization antibody titers against all SARS-Cov-2 variants (33, 34). This observation is reinforced by our finding that memory B cells induced by Wuhan-type vaccine still can bind to Omicron RBD, but just with low affinity. It is likely that more common memory B cells that can neutralize both Wuhan and Omicron strains may be selected or undergo affinity maturation.

To detect RBD-specific B cells in this study, we performed FACS analysis and ELISpot assay. FACS analysis detects the expression of surface Ig bound to RBD, while ELISpot assay can detect cells that actually secrete anti-RBD antibodies. Although this system may detect antibody-secreting cells other than IgG, there is a positive correlation between the frequency of RBD-specific memory B cells detected by these two methods. The ELISpot method has a significant advantage in the number of cells required for the measurement (10^5^ cells) compared with FACS analysis, which requires a large number of PBMCs. This is an important issue when there are limited samples of human origin.

Although much of the focus on vaccine efficacy and protective effects has focused on the role of neutralizing antibodies, T-cell responses play an important role in the resolution of infection. Memory T cells persist but gradually decline after vaccination or infection, as described by other investigators (9, 28). Even after the third booster, IFNγ-producing memory T cells recovered to the level of the second vaccination but they did not exceed these levels. Although T-cell memory generally persists even a decade after vaccination (35), it is possible that this number of memory T cells is sufficient to suppress severe injury by virus infection. Additionally, memory T cells maintain reactivity to a relatively wide range of viral antigens such as Omicron variants. This is consistent with the fact that BNT162b2 vaccine administration maintains a low protective effect against Omicron but not against severe disease (36, 37). Again, the effect of additional booster vaccination should be carefully evaluated.

An important role of helper T cells is to provide helper signals to antibody-producing B cells. In this study, we found that CXCR3^+^CCR6^-^ cTfh1 correlated positively with B cell memory, while CXCR3^-^CCR6^+^ cTFh17 showed a negative correlation. This is consistent with the importance of IFNγ from cTfh1 for the class switch to IgG. cTfh, especially cTfh1, was similarly reported to correlate positively with neutralizing antibodies in COVID-19 convalescent patients (13, 14). Additionally, in vaccinated patients, an early response of cTfh may influence the subsequent antibody response (25). In addition to these observations, we have now established a system to induce B cell differentiation by stimulating memory CD4^+^T cells *in vitro*. Because *in vitro* antibody production was dependent on stimulation with a peptide pool, it is highly likely that this response is T cell-dependent. To show T cell dependence more directly, experiments using purified T and B cells would be necessary. We found that IL-21, IFNγ, and IL-4, which are representative Tfh cytokines, were related to antibody levels in this system. In participant E014, for example, a certain level of antibodies was produced upon T-cell stimulation compared with that of other participants, and their culture results showed high IL-21, IFNγ, and IL-4 levels. We noticed that *in vitro* antibody production of samples 8 months after the second vaccination (V2_8m) was much lower than the V2_3w samples, even though memory B cell levels are higher in V2_8m samples than in V2_3w samples. Three weeks after vaccination may still stimulate innate immunity sufficient for memory cTfh cell activation which is necessary for memory B cell activation and plasma cell differentiation. Further detailed analysis will reveal the function of memory T cells in activating and amplifying B cell memory.

In this study, we also evaluated memory B and T cells induced by infection and compared them to those induced by vaccination. Although immunological memories are considered to be induced similarly by infection and vaccination (11), there seems to be a difference when we carefully compared the data; memory T cell induction was not as low with infection, but B cell memory was much lower than the after the third vaccination. This reason is not clear at present, however, presence of cross-reactivity of memory T cells to circulating “common cold” coronavirus and SARS-CoV-2 (18, 38). Unlike memory B cells, memory T cells appear to slowly decrease, and they do not show a drastic increase by booster vaccination. While other studies have shown that patients with severe disease have the strongest memory response (32), our present study showed that patients with moderate disease showed the strongest memory response. All of the patients with severe disease in our study received systemic administration of steroids, which may have inhibited the development of memory cells. However, a robust recall response in both B cell and T cell memory is expected by natural infection or by further vaccine boosting.

Finally, we examined cross-reactivity with Omicron mutants. Vaccine-induced T cell memory was as responsive to the Wuhan-type spike protein peptide as it was to the Omicron-type peptide with similar efficiency. Although the BCRs of memory B cell-bound Omicron-type RBDs, their affinity was probably severely reduced, as indicated by the decrease in fluorescence intensity of the RBD signals in the FACS analysis. However, it is possible that infection with Omicron or vaccination with Omicron-type recombinant spike protein may induce affinity maturation by helping memory Tfh cells to produce more effective antibodies.

## Acknowledgments

We would like to thank Yukiko Tokifuji, Noriko Yumoto, and Yasuko Hirata, (Keio University) for providing technical support.

## Footnotes

### Declaration of Competing Interest

The authors declare the following financial interests/personal relationships which may be considered as potential competing interests: Mitsuru Murata reports equipment, drugs, or supplies was provided by MBL Corporation. Mitsuru Murata reports equipment, drugs, or supplies was provided by Sysmex Corp. Mitsuru Murata reports equipment, drugs, or supplies was provided by Qiagen. Yoshifumi Uwamino, Mitsuru Murata, Masatoshi Wakui has patent pending to Keio University. Masaru Takeshita and Hideyuki Saya receive royalties from MBL for SARS-CoV-2 ELISA Kits.

The online version of this article contains Supplemental material.

## Abbreviations used in this article

AIM: activation induced markers
COVID-19: coronavirus disease 2019
cTfh: circulating Tfh
MFI: mean of fluorescence intensity
RBD: receptor-binding domain
RBD-ASCs: RBD-specific antibodies secreting B cells
RBD-MBCs: RBD-specific memory B cells
SARS-CoV-2: severe acute respiratory syndrome coronavirus 2
SBP: streptavidin binding peptide
Tfh: follicular helper T

