## Supplemental information for "Memory B cells and memory T cells induced by SARS-CoV-2 booster vaccination or infection show different dynamics and efficacy to the Omicron variant"

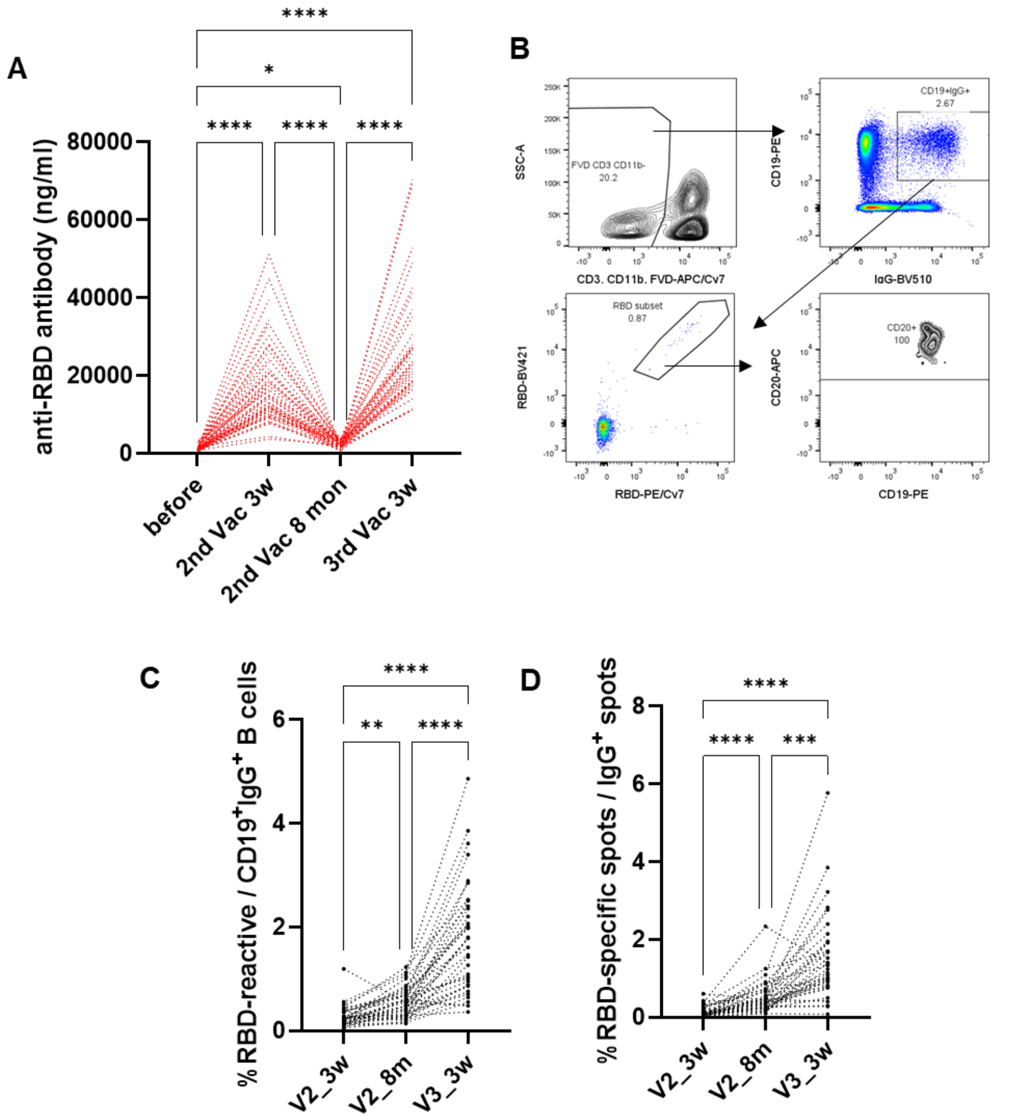


**Supplemental Figure 1.** Anti-RBD antibody concentration in plasma decrease, but the RBD-MBCs continuously increased (**A**) Each sample of Figure 1A is indicated. (**B**) Gating strategy of FACS analysis to detect RBD-MBCs is shown. (**C**) Each sample of Figure 1C. The frequencies of RBD-MBCs in each volunteer are shown. (**D**) Each sample of Figure 1F. The frequencies of RBD-ASCs in each volunteer are shown. The Kruskal-Wallis test was performed, and $p$ values were determined using Dunn's multiple comparisons test. *, $p < 0.05$; **, $p < 0.01$; ***, $p < 0.001$; ****, $p < 0.0001$.

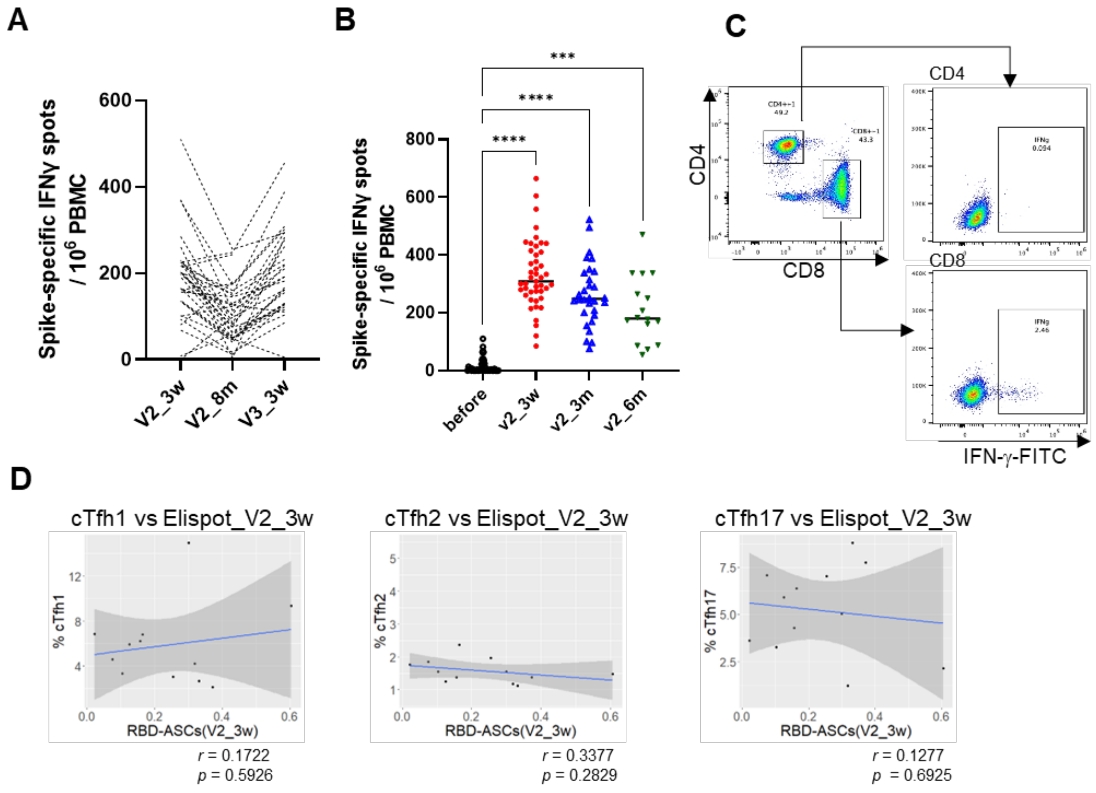


**Supplemental Figure 2.** T cell responses against RBD decrease after vaccination. (**A**) Each sample of Figure 2B. (**B**) IFNγ production at several time point after 2nd vaccination. ELISpots were detected after 2 days cultured with 15 mer oligopeptide pools. (**C**) IFNγ expression of T cells after cultured with10 mer oligopeptide pools, which activate $CD8^+$ T cell but not $CD4^+$ cells. (**D**) Pearson's correlation coefficients are calculated between each CD4 subset at V2_3w and RBD-ASCs at V2_3w. The Kruskal-Wallis test was performed, and $p$ values were determined using Dunn's multiple comparisons test. ***, $p < 0.001$; ****, $p < 0.0001$.

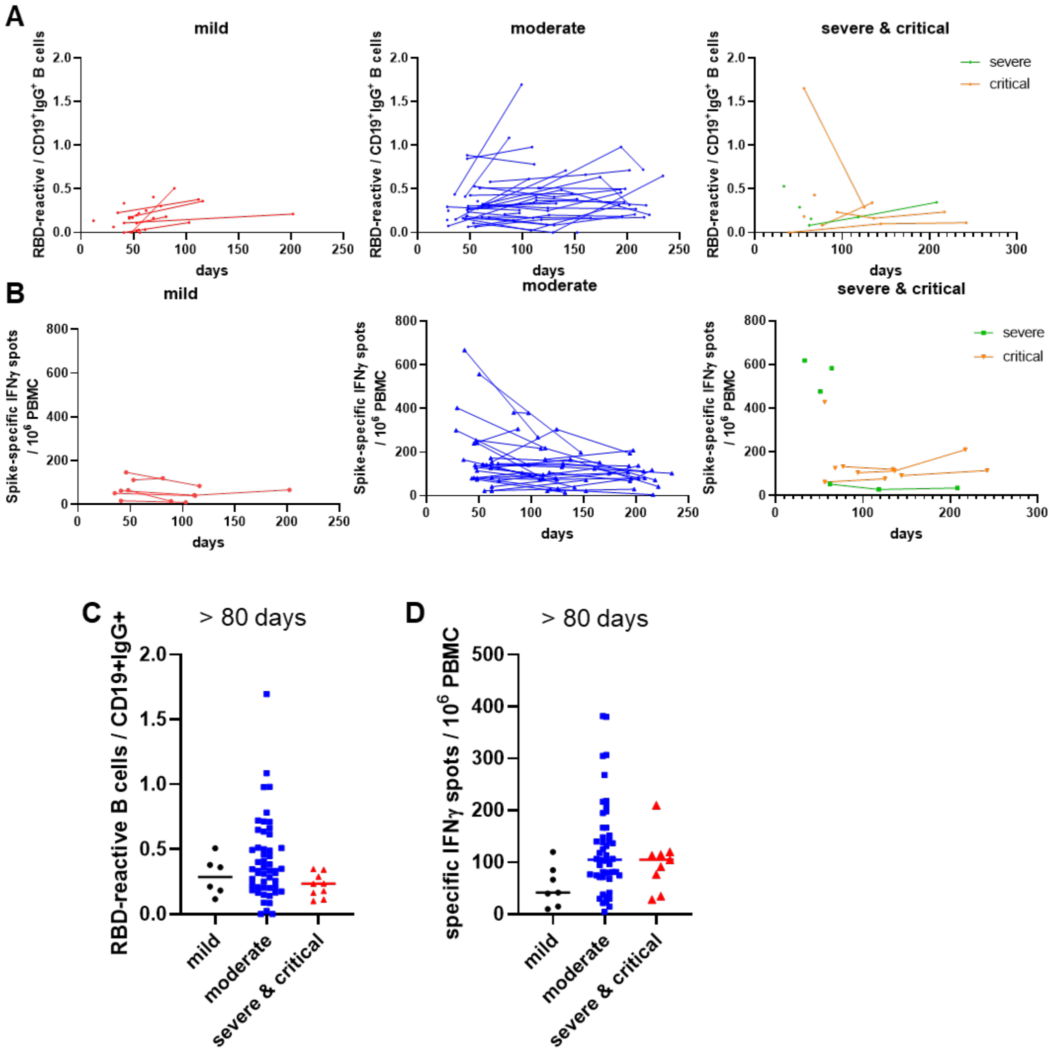


**Supplemental Figure 3.** RBD-MBCs and memory T cells in COVID-19 patients. (**A, B**) Longitudinal analyses of RBD-MBCs (**A**) and memory T cells (**B**) frequencies as a function of days after a SARS-CoV-2 PCR-positive test result are indicated in each category of COVID-19 severity. (**C, D**) The frequencies of RBD-MBCs (**C**) and memory T cells (**D**) from the COVID-19 patients over 80 days after a SARS-CoV-2 PCR-positive test result are indicated by COVID-19 severity.

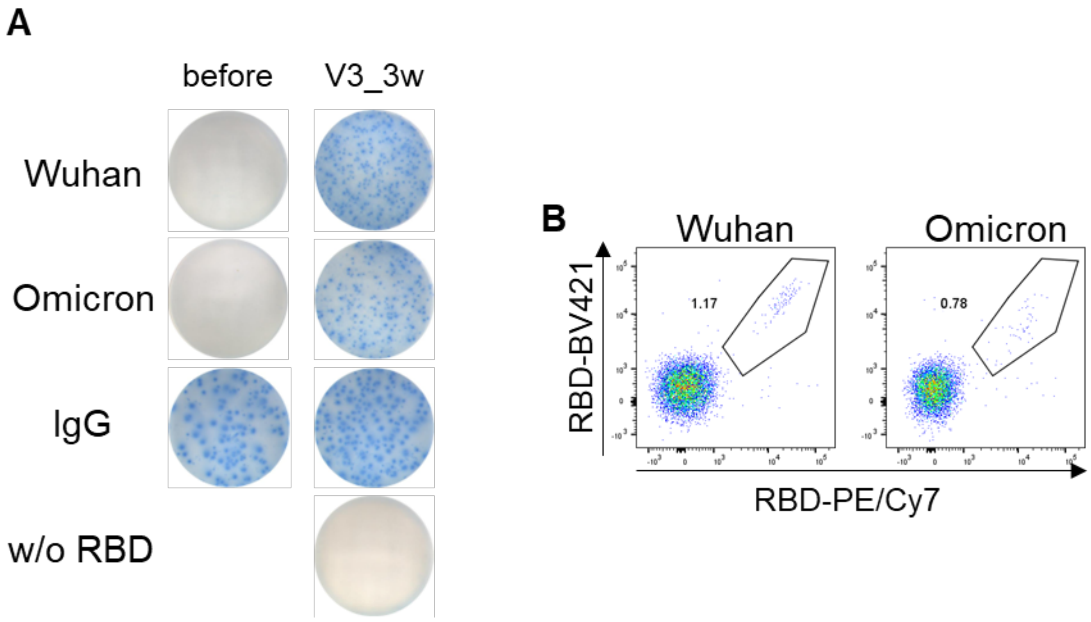


**Supplemental Figure 4**. Memory B cells are reactive to the Omicron variant. (**A**) Representative images of ELISpot assay against RBD derived from the Wuhan or the Omicron variant, or IgG before the vaccination and at 3 weeks after the third vaccination. (**B**) Representative FACS profiles of B cells reacted with RBD derived from the Wuhan or the Omicron variant are shown.
